# Enhancing Whole Slide Image Classification with Discriminative and Contrastive Learning

**DOI:** 10.1101/2024.05.07.593019

**Authors:** Peixian Liang, Hao Zheng, Hongming Li, Yuxin Gong, Yong Fan

**Author notes:** {, }.

## Abstract

Whole slide image (WSI) classification plays a crucial role in digital pathology data analysis. However, the immense size of WSIs and the absence of fine-grained sub-region labels, such as patches, pose significant challenges for accurate WSI classification. Typical classification-driven deep learning methods often struggle to generate compact image representations, which can compromise the robustness of WSI classification. In this study, we address this challenge by incorporating both discriminative and contrastive learning techniques for WSI classification. Different from the extant contrastive learning methods for WSI classification that primarily assign pseudo labels to patches based on the WSI-level labels, our approach takes a different route to directly focus on constructing positive and negative samples at the WSI-level. Specifically, we select a subset of representative and informative patches to represent WSIs and create positive and negative samples at the WSI-level, allowing us to better capture WSI-level information and increase the likelihood of effectively learning informative features. Experimental results on two datasets and ablation studies have demonstrated that our method significantly improved the WSI classification performance compared to state-of-the-art deep learning methods and enabled learning of informative features that promoted robustness of the WSI classification.

## 1 Introduction

Digital scans of pathology tissue slides, often referred to as whole slide images (WSIs), provide rich information, such as tumor microenvironments, for cancer diagnosis and treatment planning [1, 18]. While WSI classification plays an important role in addressing cancer diagnosis, it presents a significant challenge due to the gigapixel size of WSIs and the absence of pixel-level annotations.

Deep learning methods for the WSI classification task typically divide the huge WSIs into image patches and integrate the image patches for classification at the WSI-level based on features extracted from the image patches [24, 4, 3, 13]. Promising WSI classification performance has been achieved by deep learning methods with innovative graph and Transformer-based architectures that facilitate effective feature learning and patch integration for the WSI classification [5, 3, 21, 12]. Despite their promising classification performance, these classifier-driven methods face challenges in attaining compact image representations to enhance the robustness of classification accuracy in that these methods employ discriminative information alone to learn features and construct classification models, ignoring intra-class and inter-class feature variabilities [19, 30].

We aim to address this challenge and obtain compact and informative image representations for accurate WSI classification through join discriminative and contrastive learning. Contrastive learning is an effective method to learn compact feature representations by minimizing feature distances between positive samples while maximizing distances across negative samples. Existing contrastive learning methods for WSI classification can be broadly categorized into two types: self-supervised learning and weakly supervised learning. In the self-supervised methods, patches, along with their augmented or semantically similar counterparts, are regarded as positive samples while semantically dissimilar patches are considered as negative samples [11, 27, 25]. Despite the rich semantic information obtained by these methods, there exists a limitation in exploring pathology-related discriminative information since the positive and negative samples are not tied to WSI class information. In the weakly supervised learning methods, image-level labels are utilized to assign pseudo class labels to patches, forming the basis for the construction of positive and negative patch pairs [26, 22]. However, transferring the WSI-level label information to image patches may introduce class label noise and yield degraded image representations. Taking the abnormal WSI images for example: both normal and abnormal patches may coexist. Assigning abnormal pseudo labels to normal patches of the abnormal WSI can introduce extraneous noise in subsequent contrastive learning.

To overcome limitations of the extant methods, we introduce a novel framework, **DC-WSI**: **D**iscriminative and **C**ontrastive learning framework for **W**hole **S**lide **I**mage classification. Our approach employs both discriminative and contrastive learning to obtain compact and robust image representation at both the patch- and the WSI-levels for accurate WSI classification in an end-to-end multi-task WSI classification framework. Specifically, our method employs attention mechanisms to aggregate patch features for making image classification predictions. Our contrastive learning uses a set of patches to represent a WSI, with the WSI class label (discriminative) information propagated to the image patches through cross-attention, which facilitates effective learning of more robust representations of the WSIs and improves the likelihood of detecting the abnormal patches of WSIs (see Fig. 1). Such a contrastive learning strategy allows to aggregate patch features of WSIs through cross-attention for characterizing similarities between WSI images and encouraging feature learning to maximize intra-class similarity and minimize inter-class similarity.

**Fig. 1.**
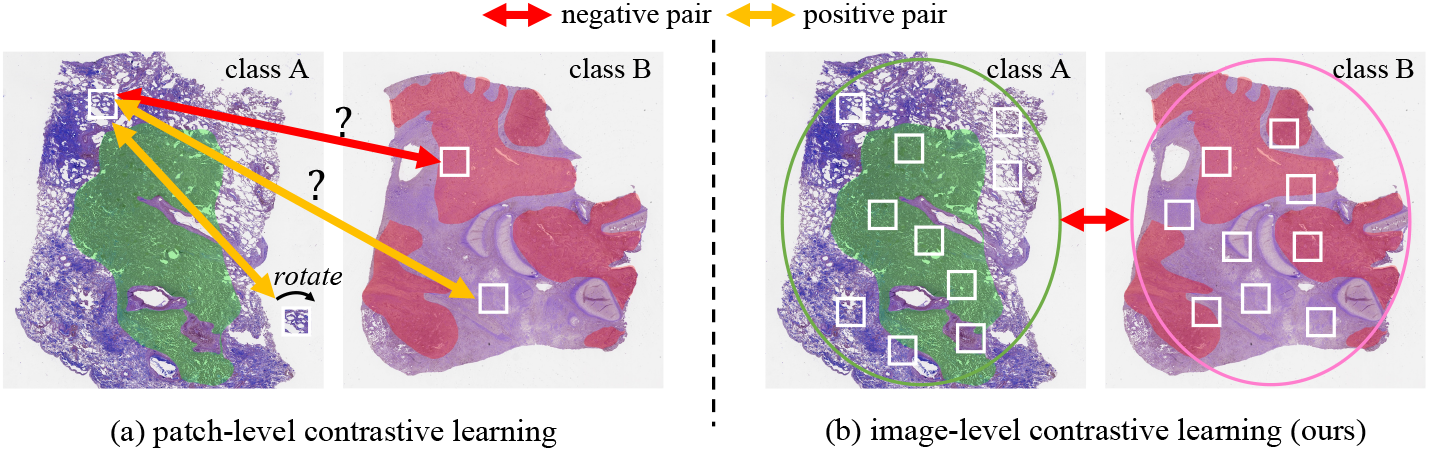
A comparison of image-level contrastive learning with patch-level contrastive learning, given two WSIs from two different cancer classes, A and B, respectively. (a) for the patch-level contrastive learning, positive and negative pairs of image patches are needed. However, the absence of patch-level label information introduces potential noise in both positive and negative pairs, while self-supervised contrast learning cannot utilize class label information. (b) for the image-level constrastive learning, positive and negative samples are defined based on the class label information of WSIs, and such information is propagated to the image patches for enhancing feature learning with our proposed method. By treating a set of patches as the basic unit, it allows to learn a more comprehensive representation of WSIs and increases the likelihood of capturing cancerous regions. Cancer areas are depicted by the green and red colors in the corresponding images.

Our contributions are three-folds: ***1)*** We propose a new discriminative and contrastive learning framework to learn compact and robustness image representations for accurate WSI classification; ***2)*** We propose a new WSI-level contrastive learning method with a set of patches to refrain from using patch-level pseudo labels and thus mitigate the label noise in the learning process; and ***3)*** We design a deep learning model with joint discriminative and contrastive learning to improve the WSI classification performance.

## 2 Method

### 2.1 Problem Definition

Given a set of WSIs 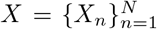, which is split into two parts: training set 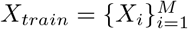 and testing set 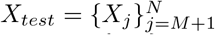. Each training WSI *X*_*i*_ ∈ *X*_*train*_ has its binary image class label *Y*_*i*_ ∈ {0, 1} representing normal/abnormal or different disease types. Our goal is to predict class label *Y*_*j*_ ∈ {0, 1} for each test image *X*_*j*_ ∈ *X*_*test*_.

### 2.2 Method Overview

Our method is schematically illustrated in Fig. 2, consisting of two parts: *Patch Selection* and *DC-WSI Model*. (1) *Patch Selection*: Selecting a subset of informative and representative patches to represent each of the WSIs to facilitate computationally efficient WSI classification. Specifically, given a WSI *X*_*i*_, we divide it into *m* non-overlapping patches (*m* can vary for different WSIs). A foundation model is then applied to encode each patch into a fixed dimensional vector to capture the semantic information of the patch. All vectors within a WSI are then input into a clustering method to group the patches into *k* clusters. Subsequently, *q* patches are randomly selected from each cluster, forming a set *P*_*i*_ ={ *p*_*i*,1_, *p*_*i*,2_, …, *p*_*i,b*_}, where *b* is the number of selected patches. *P*_*i*_ is used for further training. (2) *DC-WSI Model* : We construct an end-to-end dual discriminative and contrastive learning model to predict class labels of WSIs. For the contrastive learning, we sample a pair of WSIs, *X*_1_, *X*_2_ ∈ *X*_*train*_ from the training set. If they belong to the same class, their corresponding patch set *P*_1_ and *P*_2_ are considered as a positive pair; otherwise, it is a negative pair. An encoder is then applied to each patch *p*_*i,j*_, *i* ∈ { 1, 2 }to produce a fixeddimensional vector *f*_*i,j*_ that captures disease-aware information. After encoding, we transform patch set *P*_*i*_ into the encoding space 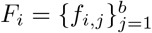. Subsequently, a self-attention module aggregates features *F*_*i*_ to make a classification prediction for *X*_*i*_. Additionally, a contrastive learning module using cross-attention layers to characterize similarity between *X*_1_ and *X*_2_ with a similarity score *sim*_1,2_, based on their patch features *F*_1_ and *F*_2_. The contrastive learning loss is designed to encourage positive samples to be similar to each other while pushing negative samples apart.

**Fig. 2.**
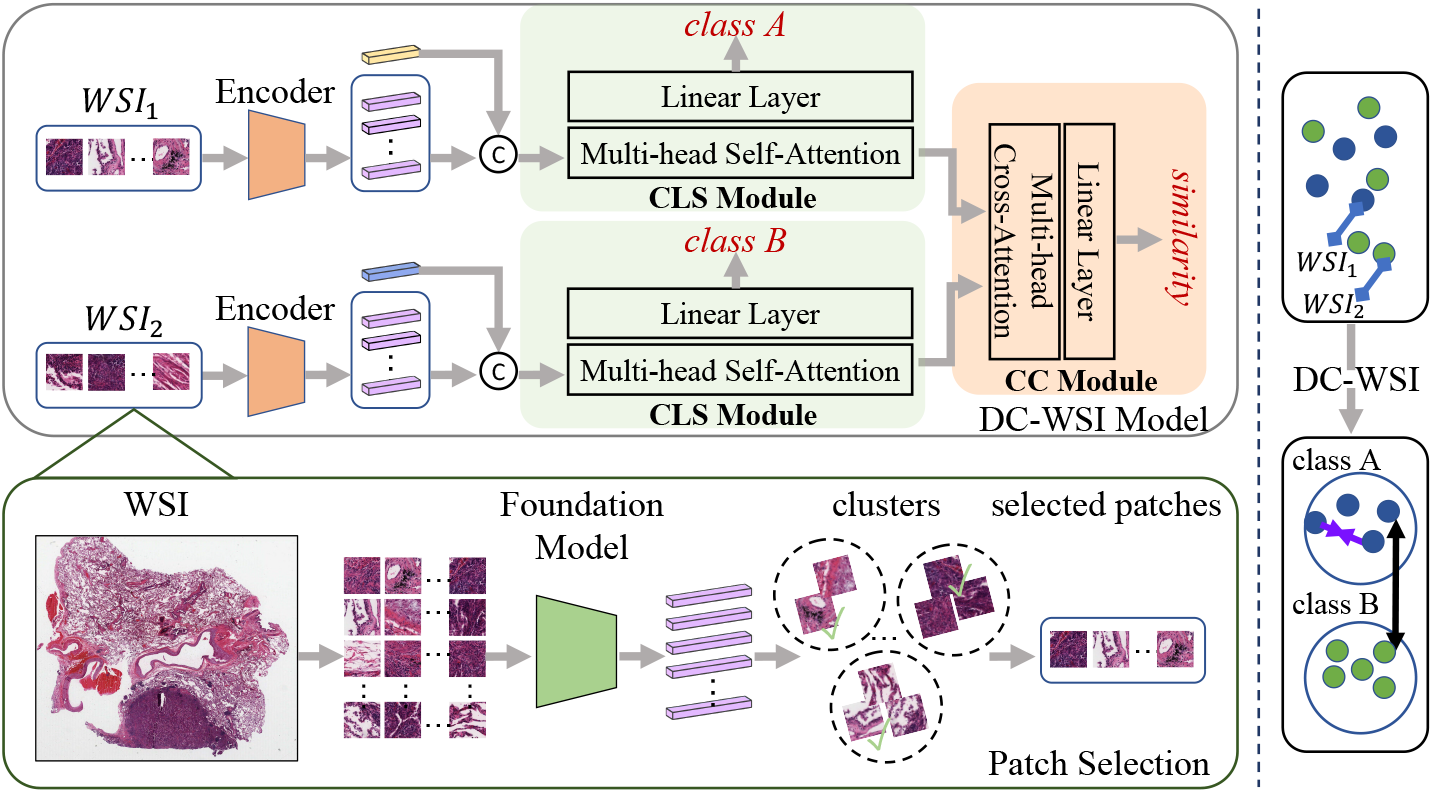
An overview of our proposed DC-WSI method. Two WSIs are sampled from the training set for demonstration. Given each WSI representation (i.e., a set of selected patches), an encoder is applied to extract patch features. The classification (CLS) module aggregates intra-image features to predict class labels, and the contrastive learning (CC) module facilitates effective learning of informative features that maximize intraclass similarity and minimize inter-class similarity.

### 2.3 Representative Patch Selection

We apply SAM [10] as the feature extractor to obtain patch features for subsequent clustering. SAM is a robust foundation model capable of extracting dataset-agnostic semantic information. Specifically, we employ the SAM encoder to transform patches into fixed-size one-dimensional vectors. Once all patch features within a WSI are obtained, we apply the K-means clustering method [16] to categorize the corresponding patches into *k* clusters. For each cluster, we randomly sample *q* patches. The collection of selected patches form a patch set *P*_*i*_ = {*p*_*i*,1_, *p*_*i*,2_, …, *p*_*i,b*_}, where *b <*= *m. P*_*i*_ denotes a WSI *X*_*i*_ to be used for feature learning and WSI classification.

### 2.4 DC-WSI Model

#### Encoder

We employ ResNet18 [7] as an encoder backbone, excluding its last three layers. Given a WSI with informative and representative patches 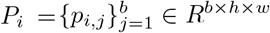, where *h × w* is the patch size, it is used as an input to the encoder for learning patch features: 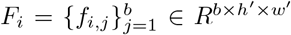, where *h*^*′*^ *× w*^*′*^ represents feature map size. These patch features are then flattened to produce 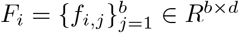, where *d* = *h*^*′*^ *× w*^*′*^. *F*_*i*_ is to be optimized through training by minimizing a WSI classification loss and a contrastive learning loss in an end-to-end training process.

#### Classification Module

The classification (CLS) module aims to predict a class label of *X*_*i*_ based on its patch features *F*_*i*_. Specifically, following the setting of ViT [6], we add a learnable class token *C* ∈ *R*^1*×d*^ into the patch features *F*_*i*_ to learn a set of features 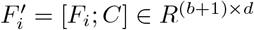. Then, 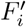 is fed into a Multi-head Self-Attention module. Specifically, 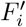 goes through a *multi-head attention* [23] layer, which yields query, key, and value vectors: 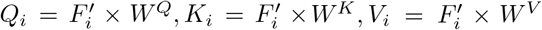, where *W*^*Q*^, *W*^*K*^, *W*^*V*^ ∈*R*^*d×d*^ are parameter matrices. Finally, for each head *j* an attention output is computed as:

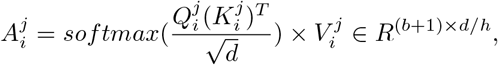

where 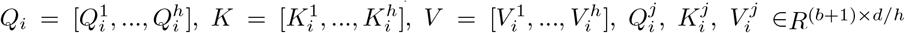, and *h* is the number of heads.

The attention outputs from all heads are concatenated to form a feature set 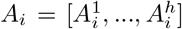. The class token 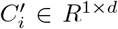 is taken from *A*_*i*_ and passed through a linear layer to generate a prediction of the class probability, denoted as *q*_*i*_ ∈ *R*. The prediction is supervised by the image label *Y*_*i*_, and the classification loss *L*_*cls*_ is computed as a binary cross-entropy loss:

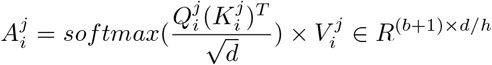

#### Contrastive Learning Module

The Contrastive Learning (CL) module operates on pairs of images *X*_*i*_ and *X*_*j*_ with corresponding class labels of *Y*_*i*_ and *Y*_*j*_ respectively, sampled from the training set *X*_*train*_. The objective is to optimize the image features by maximize intra-class similarity and minimize inter-class similarity. Positive pairs contain images with the same class label, i.e., *Y*_*i*_ = *Y*_*j*_, while negative pairs contain images from different classes.

Specifically, for a pair of images with features of *A*_*i*_ and *A*_*j*_, a *multi-head* attention is used to obtain cross-attention between them for computing their similarity, *sim*_*i,j*_ ∈ [0, 1], between their class tokens *C*_*i*_ and *C*_*j*_ with a linear layer.

During each training iteration, *Z* pairs of images are sampled, with *Z/*2 are negative samples, and *Z/*2 are positive samples. The similarity scores of all positive pairs are added to derive *SIM*_*pos*_, while the similarity scores of all negative pairs are added to derive *SIM*_*neg*_. The contrastive learning loss *L*_*cc*_ is calculated using the maximum-margin classification loss:

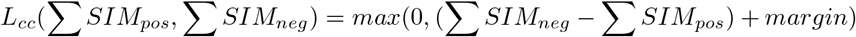

The final loss is the sum of the classification loss *L*_*cls*_ and the contrastive learning loss *L*_*cc*_. The model is trained end-to-end. During inference, only the classification branch is utilized to predict class labels for input WSIs.

## 3. Experiments

We conduct experiments on two datasets: TCGA-Lung and TCGA-ESCA to demonstrate the effectiveness of the proposed WSI-CL method. Additionally, we perform ablation studies to demonstrate the effectiveness of key components in our WSI-CL method.

### Datasets *1)*

TCGA-Lung is a public dataset from National Cancer Institute Data Portal [2]. It includes two types of lung cancer, i.e., Lung Squamous Cell Carcinoma (TCGA-LUSC) and Lung Adenocarcinoma (TCGA-LUAD). A total of 1042 diagnostic WSIs were collected and randomly divided into training and testing sets with a ratio of 0.8:0.2. The training set contained 409 TCGA-LUSC and 424 TCGA-LUAD, and the test set contained 103 TCGA-LUSC and 106 TCGA-LUAD. ***2)*** TCGA-ESCA is a dataset from National Cancer Institute Data Portal [2]. A total of 149 diagnostic slides are collected which contains two types of Oesophageal Carcinoma, i.e., Squamous cell carcinoma (SCC) and Adenocarcinoma (AD). We randomly split it into training and testing sets with a ratio 0.8:0.2. The training set contained 68 SCC and 49 AD, and the testing set contained 16 SCC and 16 AD.

### Evaluation Metrics

We used the standard WSI classification evaluation metrics, including *Accuracy* [28] and area under the curve (*AUC*) score [29]. Specifically, 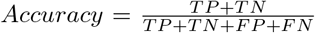, where TP=True positive, TN=True negative, FP=False positive, FN=False negative.

### Implementation Details

In our experimental setup, WSIs were partitioned into patches of size 224x224 at 10X magnification. For the patch selection, we used following parameters: *k* = 8 and *q* = 50. In the contrastive learning module, we configured *Z* = 6. The learning rate was set to 0.0002, and the optimization was performed using the Adam optimizer [9]. The model was implemented using PyTorch [17].

### Comparison Methods

We compared our method with an array of state-of-the-art (SOTA) WSI classification methods. Particularly, ABMIL [8] is a multi instance learning (MIL) framework to aggregate instance features through attention for final bag-level prediction, TransMIL [20] is a Transformer based WSI classification framework to aggregate patch features by attention mechanism, GTP [31] is a Transformer-Graph based WSI classification framework with a patch-level contrastive learning to learn patch features, and CLAM [14] is a clustering-constrained attention MIL approach with a patch clustering loss to impose constraints and refine patch features in the WSI classification process.

### 3.1 WSI Classification Results and ablation studies

Table 1 shows WSI classification comparison results on TCGA-Lung and TCGA-ESCA datasets. Firstly, our method (*DC-WSI (PS+L*_*cls*_ + *L*_*cc*_)) obtained the overall best WSI classification performance among all methods under comparison. Particularly, our method achieved substantial improvement on the TCGA-ESCA dataset, with a 6.30% increase in *Accuracy* and a 1.6% increase in *AUC* compared to the second-best method. On the TCGA-Lung dataset, our approach achieved a 0.4% increase in *Accuracy* over the second-best method. These results demonstrated the effectiveness of our method in extracting discriminative features and aggregating them for accurate WSI classification predictions.

**Table 1.** WSI classification comparison results on TCGA-Lung and TCGA-ESCA datasets. RS denotes random sampling, PS denotes proposed patch selection. The **bold** score represents the best performance on the corresponding dataset.

|  | TCGA-Lung |  | TCGA-ESCA |  |
| --- | --- | --- | --- | --- |
| | Accuracy (%) ↑ | AUC (%) ↑ | Accuracy (%) ↑ | AUC (%) ↑ |
| ABMIL [8] | 0.785 | 0.866 | 0.750 | 0.846 |
| TransMIL [20] | 0.823 | 0.905 | 0.781 | 0.922 |
| GTP [31] | 0.876 | **0.956** | 0.875 | 0.949 |
| CLAM [14] | 0.876 | 0.952 | 0.875 | 0.941 |
| DC-WSI (RS+$L_{cls}$) | 0.828 | 0.905 | 0.875 | 0.933 |
| DC-WSI (PS+$L_{cls}$) | 0.856 | 0.916 | 0.906 | 0.938 |
| DC-WSI (PS+$L_{cls}$ + $L_{cc}$) | **0.880** | 0.940 | **0.938** | **0.965** |

Secondly, ablation studies demonstrated the effectiveness of key components of our method. *DC-WSI (RS+L*_*cls*_) denotes the ablation study of representative patch selection in Section 2.3. Instead of using our proposed patch selection strategy, *DC-WSI (RS+L*_*cls*_) randomly selected the same amount of patches with representative patch selection for further training. The results in Table 1 showed that our patch selection strategy *DC-WSI (PS+L*_*cls*_) outperformed *DC-WSI (RS+L*_*cls*_), indicating that our patch selection strategy can capture more informative patches, leading to more accurate WSI classification.

*DC-WSI (PS+L*_*cls*_) denotes the contrastive learning ablation study. Instead of using both discriminative and contrastive learning modules, *DC-WSI (PS+L*_*cls*_) only used the classification module with *L*_*cls*_ loss. Table 1 shows results. The performance degradation of *DC-WSI (PS+L*_*cls*_) compared to *DC-WSI (PS+L*_*cls*_ + *L*_*cc*_) demonstrated the effectiveness of the contrastive learning component in improving the classification performance.

Fig. 3 shows a t-SNE [15] visualization comparison of image representations (i.e., class tokens *C*_*i*_) obtained from *DC-WSI (PS+L*_*cls*_) and *DC-WSI (PS+L*_*cls*_ + *L*_*cc*_) models respectively, further demonstrating that the contrastive learning module can help learn more informative features that maximized the intra-class similarity and minimized the inter-class similarity, compared with the classification model with the discriminative learning alone.

**Fig. 3.**
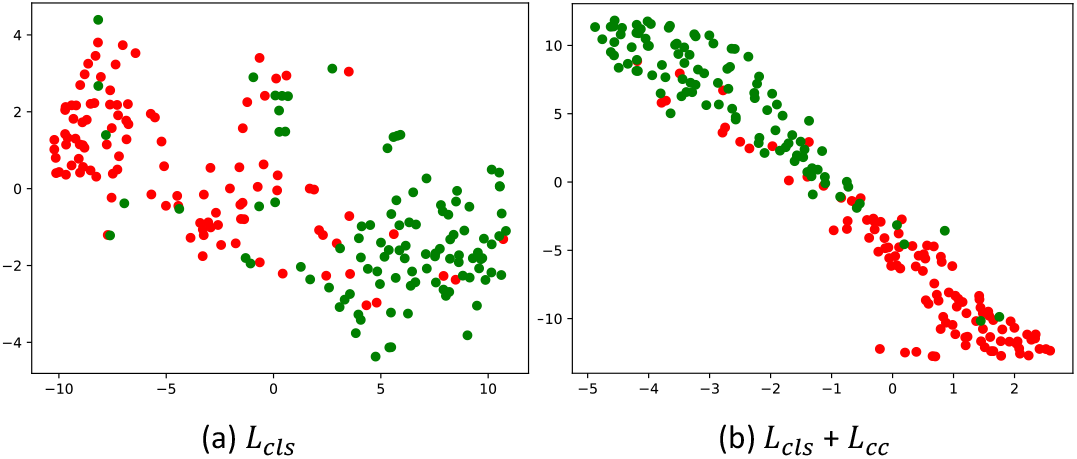
t-SNE visualization of image features (class tokens) of WSIs learned by (a) a classification model (*L*_*cls*_) and (b) a classification+contrastive model (*L*_*cls*_ + *L*_*cc*_) on TCGA-Lung test set. Different colors represent samples in different classes.

## 4. Conclusion

We develop a new discriminative and contrastive learning framework for WSI classification. Experimental results on two WSIs datasets and ablation studies have demonstrated that the proposed method can learn discriminative features that improved WSI classification performance, maximized intra-class similarity, and minimized inter-class similarity. Specifically, our method selects a subset of informative and representative patches as the basic unit of WSIs, while positive and negative samples are directly constructed at the WSI-level for the contrastive learning. Compared to the extant patch-level-based contrastive learning for the WSI classification, utilization of a set of patches as a basic unit not only facilitates effective learning of robust features from the WSIs but also improves classification performance. Our method can be further improved by incorporating the patch selection in the end-to-end learning, though our current strategy offers the flexibility to use different clustering algorithms to select representative patches.

